# Expression, purification, and characterization of diacylated Lipo-YcjN from *Escherichia coli*

**DOI:** 10.1101/2024.09.05.611266

**Authors:** Matthew A. Treviño, Kofi Amankwah, Daniel Fernandez, Scott Weston, Claire J. Stewart, Jaime Morales Gallardo, Mona Shahgholi, Naima G. Sharaf

## Abstract

YcjN is a putative substrate-binding protein expressed from a cluster of genes involved in carbohydrate import and metabolism in *Escherichia coli*. Here, we determine the crystal structure of YcjN to a resolution of 1.95 Å, revealing that its three-dimensional structure is similar to substrate binding proteins in subcluster D-I, which includes the well-characterized maltose binding protein (MBP). Furthermore, we found that recombinant overexpression of YcjN results in the formation of a lipidated form of YcjN that is posttranslationally diacylated at cysteine 21. Comparisons of size-exclusion chromatography profiles and dynamic light scattering measurements of lipidated and non-lipidated YcjN proteins suggest that lipidated YcjN aggregates in solution via its lipid moiety. Additionally, bioinformatic analysis indicates that YcjN-like proteins may exist in both Bacteria and Archaea, potentially in both lipidated and non-lipidated forms. Together, our results provide a better understanding of the aggregation properties of recombinantly expressed bacterial lipoproteins in solution and establish a foundation for future studies that aim to elucidate the role of these proteins in bacterial physiology.

## Introduction

ATP-binding cassette (ABC) transporter systems are present in all organisms and mediate the transport of molecules across cellular membranes.^1^ These systems share a conserved architecture consisting of transmembrane domains (TMDs) that act as a channel for ligand transport and nucleotide binding domains (NBDs) that hydrolyze ATP. Some ABC transporters also associate with substrate binding proteins (SBPs) that bind molecules and deliver them to their cognate ABC transporter.^2^ To create a functional complex, SBPs and ABC transporters must be delivered to their appropriate subcellular locations. In diderm bacteria, which contain an inner membrane (IM) and an outer membrane (OM), SBPs are secreted through the IM into the periplasm, and ABC transporters are embedded within the IM. The appropriate assembly of an ABC transporter ensures that TMDs and NBDs can access periplasmic SBPs and cytoplasmic ATP, respectively (Figure 1A).

**Figure 1:**
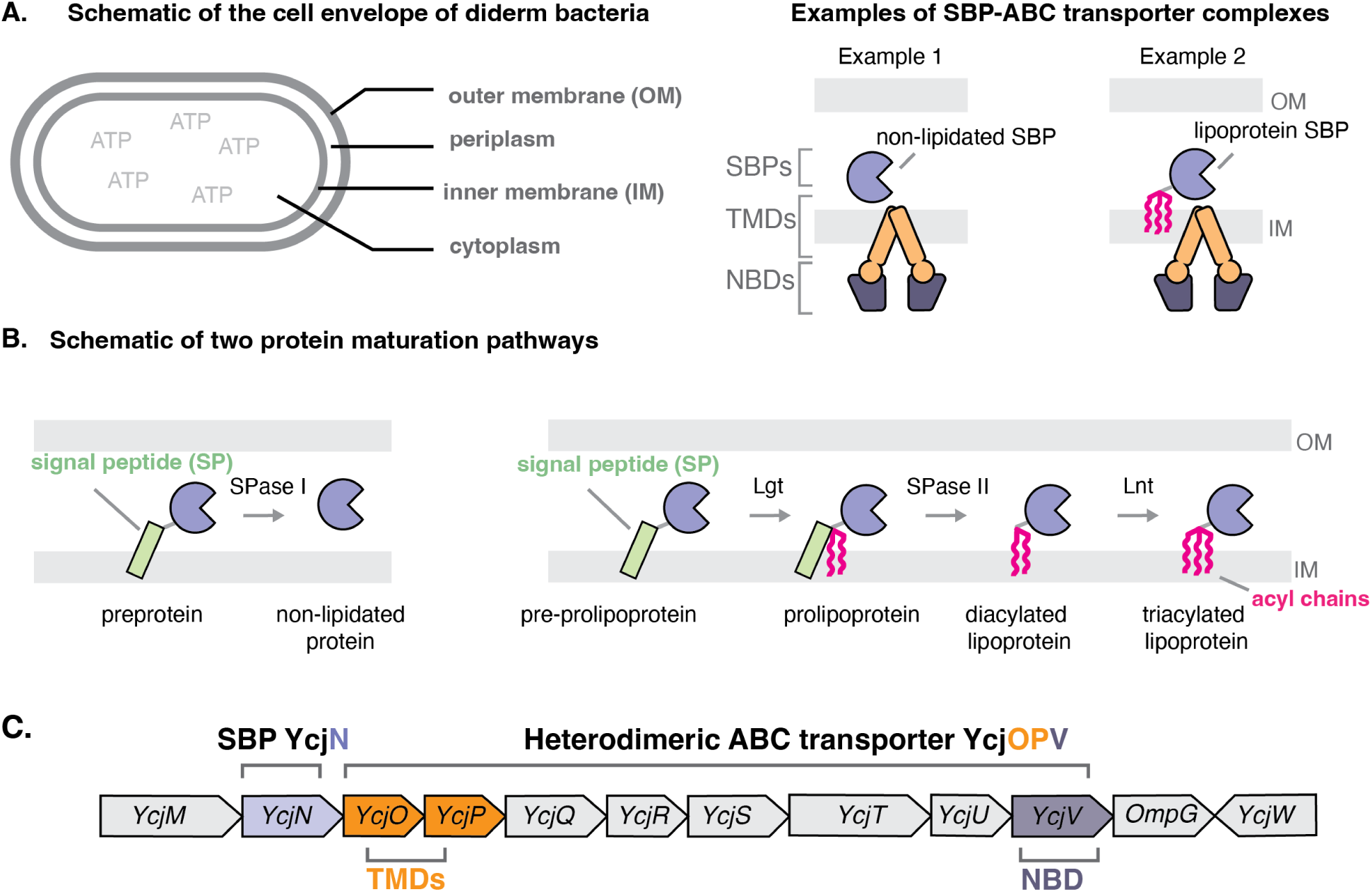
Subcellular localization and posttranslational modification pathways of SBPs. A. Diagram of the cell envelope (left panel) and organization of ABC transporter systems in complex with a soluble SBP and a lipoprotein SBP(right panel), respectively. B. Non-lipidated protein maturation pathway (left panel) and lipoprotein maturation pathway (right panel). C. Representation of the *ycj* gene cluster in *E. coli*. The protein encoding genes for the SBP, TMDs, and NBD are colored in light blue, orange, and dark blue, respectively.

Since SBPs are produced in the cytoplasm, they must first traverse the IM as immature preproteins. While attached to the IM, these proteins can undergo a series of posttranslational modifications. Studies on protein maturation have revealed that this process depends on distinct protein segments, specifically the N-terminal signal peptide (SP) (Figure 1B, green rectangles). For example, preproteins destined to mature into untethered secreted soluble proteins (referred to here as non-lipidated proteins) contain an SP recognized by type I signal peptidase (SPase I) (Figure 1B, left panel). This enzyme cleaves the C-terminal region of the SP from the IM-tethered preprotein, releasing the mature protein into the periplasm.^3^ Alternatively, preproteins that mature into lipoproteins contain an SP recognized by phosphatidylglycerol:prolipoprotein diacylglyceryl transferase (Lgt) (Figure 1B, right panel). This enzyme adds a diacylglyceryl group from phospholipids to the sulfhydryl group of the invariant cysteine within the C-terminal region of the SP. Type II signal peptidase (SPase II) then cleaves the SP so that the S-diacylglyceryl-cysteine becomes the new N-terminus. A third enzyme, apolipoprotein N-acyltransferase (Lnt), then acylates the amino group of cysteine to produce a triacylated N-acyl S-diacylglyceryl-lipoprotein (Figure 1B, right panel).^4^ To date, most SBPs characterized in diderm bacteria mature into non-lipidated SBPs. However, some SBPs have been experimentally determined to be lipoproteins.^5–7^

SBPs associate with ABC transporters to scavenge nutrients from their surroundings, such as sugars, trace metals, and amino acids.^8–10^ Despite their low sequence similarity, SBPs have a highly conserved three-dimensional fold that consists of two *α*/*β* domains and a hinge region with one to three segments. Ligand binding causes SBP conformational rearrangements that result in the formation of a ligand binding pocket at the interface between these domains. In this pocket, the SBP and the ligand form an extensive network of hydrophobic interactions and hydrogen-bond contacts, which influence ligand binding specificity. A previous study grouped SBPs into seven clusters (A-G) based on structural similarity. While there is some correlation between the cluster and the ligand specificity, SBPs within the same cluster can bind to a wide range of ligands.^11^ This observation suggests that SBP ligand specificity cannot be determined by structural comparison alone and should be verified experimentally.

YcjN is a poorly characterized putative SBP in *Escherichia coli* (*E. coli*), whose protein-encoding gene is located within a cluster of 12 genes. ^12^ In this cluster, three genes are predicted to encode protein components of an ABC transporter, including two TMDs (YcjO and YcjP) and an NBD (YcjV). Additionally, this operon also encodes two sugar dehydrogenases (YcjS and YcjQ), two phosphorylases (YcjT and YcjM), a *β*-phosphoglycomutase (YcjU), an epimerase/isomerase (YcjR), a predicted LacI-type repressor (YcjW), and an outer-membrane porin believed to be involved in the nonspecific import of oligosaccharides (OmpG) (Figure 1C). Furthermore, previous studies using radioactive palmitate labeling have shown that YcjN is a lipoprotein.^6^ However, YcjN’s site of lipidation, its molecular structure, and its ligand binding specificity remain unknown.

Here, we used various biophysical tools to characterize *E. coli* YcjN. Specifically, using liquid chromatography mass spectrometry (LC-MS) of recombinantly overexpressed YcjN, we show that it is heterogeneously diacylated at cysteine 21. A comparison to its non-lipidated protein form (ΔYcjN) using dynamic light scattering (DLS) also reveals that Lipo-YcjN aggregates via its lipid moiety in solution. We also report the first crystal structure of the non-lipidated YcjN form (ΔYcjN) to a resolution of 1.95 Å. A structural comparison of YcjN to previously characterized SBPs reveals that YcjN can be classified into subcluster D-I and is structurally similar to the well-characterized maltose binding protein (MBP). Next, we investigated the taxonomic distribution of YcjN-like proteins using bioinformatic tools. Our analyses suggest that YcjN-like proteins are present in both Bacteria and Archaea. Together, our work provides a more detailed understanding of the aggregation properties of Lipo-YcjN in solution and establishes a foundation for future studies aimed at elucidating the role of YcjN in bacterial physiology.

## Results

### Recombinantly overexpressed *E. coli* YcjN is a lipoprotein heterogeneously lipidated at cysteine 21

The SPs of bacterial lipoproteins often contain a consensus sequence [LVI][ASTVI][GAS]-[C], known as the lipobox motif, which is located within the first 40 N-terminal residues.^13^ A visual inspection of YcjN’s SP revealed two lipobox motifs, [L_12_VSC_15_] and [I_17_SGC_21_], each containing a cysteine residue with the potential to be modified with lipids (cysteine 15 and 21) (Figure 2A, pink triangles). To determine whether cysteine 15 or cysteine 21 is lipidated, we recombinantly overexpressed YcjN in *E. coli* with its native N-terminal SP and a C-terminal decahistidine tag to aid in protein purification (SP-YcjN) (Figure 2A). The full-length SP-YcjN protein was then purified using affinity followed by size-exclusion chromatography (SEC) in the presence of the detergent 2,2-didecylpropane-1,3-bis-*β*-D-maltopyranoside (LMNG) (Figure 2B, dark blue trace). The SEC elution profile displayed major and minor peaks with elution volumes of 62 mL and 84 mL, respectively (Figure 2B, dark blue star and square, respectively). SDS-PAGE analysis of peak fractions revealed prominent bands close to the theoretical molecular weight of SP-YcjN (48,180 Da) (Figure S1.A). These data suggest that we successfully expressed and purified recombinant SP-YcjN. Furthermore, SEC and SDS-PAGE analysis reveal that there are two species of SP-YcjN with similar molecular weights but with different elution volumes.

**Figure 2:**
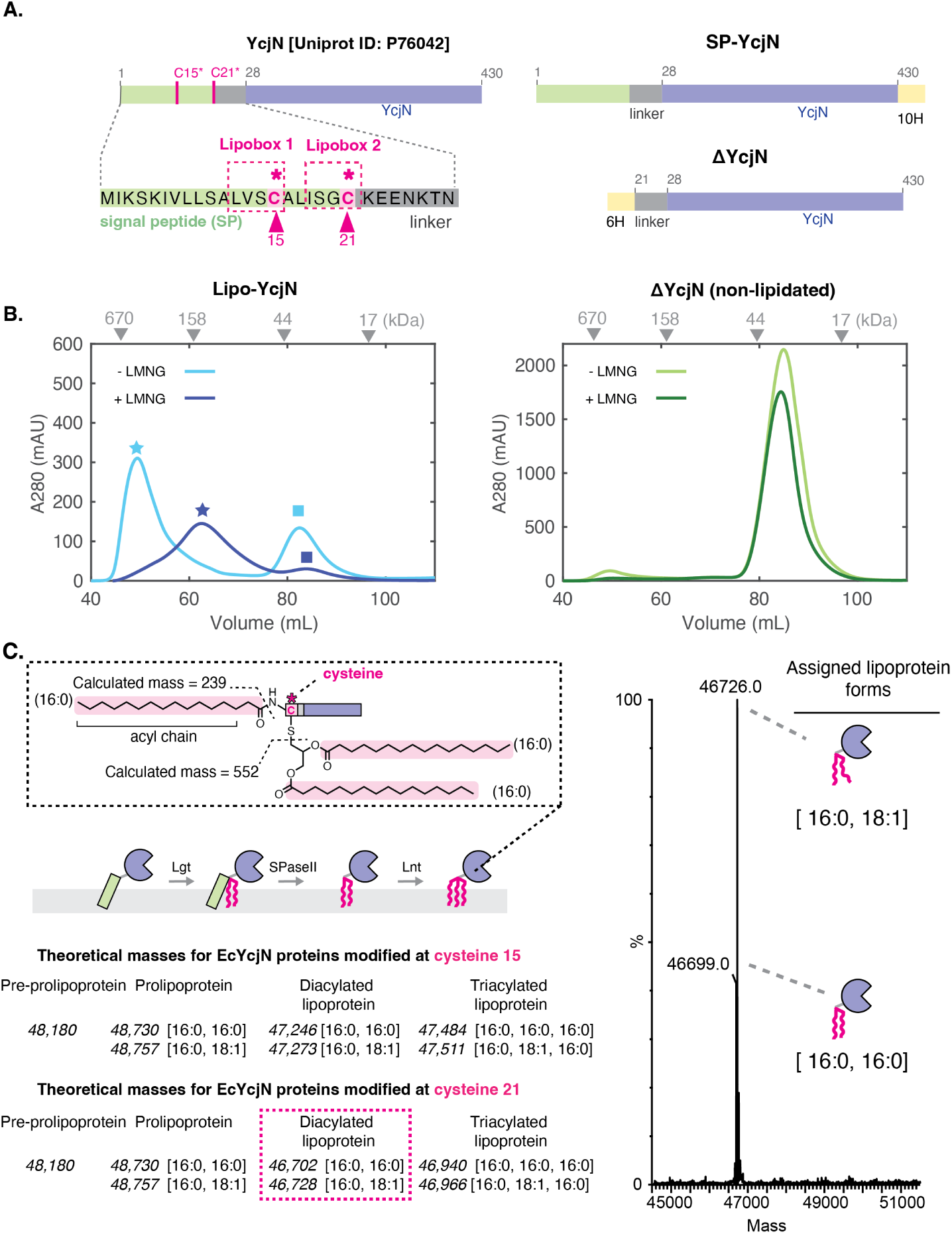
Purification and mass spectrometry analysis of YcjN proteins. A. Diagram of the amino acid sequences of YcjN proteins. Full-length YcjN contains two distinct lipoboxes (pink dashed boxes) within its SP (green rectangle). Each lipobox contains a cysteine residue (15 and 21, pink triangles) B. SEC elution profiles of Lipo-YcjN purified in the presence and absence of LMNG are shown in dark and light blue, respectively. SEC elution profiles of ΔYcjN purified in the presence and absence of LMNG are shown in dark and light green, respectively. Peak elution volumes of gel filtration standards are indicated by grey triangles. C. The theoretical masses of YcjN proteins modified at cysteine 15 (top) or cysteine 21 (bottom) are calculated, assuming that triacylation occurs via the canonical triacylation pathway due to the sequential action of three enzymes (Lgt, SPase II, and Lnt). Grey dashed box insert contains illustration of acyl chains attached to a C-terminal cysteine and the average calculated acyl chain masses of a lipoprotein with acyl chain composition [16:0, 16:0, 16:0]. Mass spectrum of the peak fraction (elution volume of 62 mL, dark blue star) of Lipo-YcjN purified in the presence of LMNG. The molecular masses of the major species correspond within 3 Da of the predicted masses of two diacylated YcjN proteins modified at residue 21, one protein with an acyl chain composition [16:0,16:0] and another with [16:0,18:1] (right panel). Assigned lipoprotein forms are shown as illustrations on the right.

To determine whether cysteine 15 or cysteine 21 is lipidated, we used LC-MS for intact protein analysis of the major peak fraction of SP-YcjN purified in the presence of LMNG (Figure 2B, dark blue star marker). This method was selected because of the distinct theoretical mass differences between YcjN lipoprotein forms. For example, the mass difference between the triacylated YcjN lipoproteins at cysteine 15 and cysteine 21 is approximately 545 Da (Figure 2C, left panel). Mass spectrometry analysis showed deconvoluted mass values of 46,699 Da and 46,726 Da (Figure 2C), aligning closely with the theoretical masses of YcjN lipoproteins modified at cysteine 21 with diacyl chain compositions of [16:0, 16:0] (46,702 Da) and [16:0, 18:1] (46,728 Da), respectively. The numbers in square brackets indicate the total number of carbons and double bonds in each acyl chain, respectively. The observed mass values do not match the theoretical masses of YcjN proteins lipidated at cysteine 15, which would be approximately 47,500 Da. These findings suggest that recombinantly overexpressed lipidated YcjN is heterogeneously diacylated at cysteine 21 (referred to as Lipo-YcjN).

### Lipo-YcjN can be purified with and without LMNG

In our previous study, we used detergents for the purification of *N. meningitidis* lipoprotein MetQ^5^. We reasoned that detergents would facilitate the extraction of this lipoprotein from cellular membranes. However, it was unknown whether detergent is essential for the purification of all bacterial lipoproteins. To determine whether diacylated Lipo-YcjN can be purified in the absence of detergent, we repeated the purification of Lipo-YcjN in detergent-free conditions. A representative SEC profile of Lipo-YcjN purified in the absence of detergent is shown in Figure 2B (light blue trace). This profile exhibited a major peak with an elution volume of 49 mL (Figure 2B, light blue star) and a minor peak with an elution volume of 82 mL (Figure 2B, light blue square). An SDS-PAGE analysis of the peak fractions revealed prominent bands close to the theoretical molecular weight of Lipo-YcjN (46,702 Da) (Figure S1.B). These data suggest that detergent is not essential for the purification of recombinantly overexpressed diacylated Lipo-YcjN using chromatography techniques.

### Lipo-YcjN aggregates via its lipid moiety

The SEC profile of Lipo-YcjN purified in the absence of LMNG reveals that its major peak elution volume is lower than anticipated (Figure 2B, light blue star). Specifically, we expect proteins with masses similar to monomeric Lipo-YcjN (46,702 Da, theoretical molecular weight of YcjN heterogenously lipidated at residue 21) to elute at approximately 85 mL, as determined by the peak elution volumes of gel filtration standards (Figure 2B, grey triangles). However, Lipo-YcjN elutes at 49 mL, an elution volume similar to that of a gel filtration protein standard with a molecular weight close to 670,000 Da (Figure 2B, light blue star). These results suggest that Lipo-YcjN aggregates in solution.

To determine whether Lipo-YcjN aggregation is mediated primarily by its globular domain or lipid moiety, we designed a construct with an SP deletion (ΔYcjN, residues 1-21 deleted) (Figure 2A). In this experiment, if aggregation is driven primarily by the protein’s globular domain, we would anticipate that both ΔYcjN and Lipo-YcjN would elute at similar volumes. In other words, aggregation is not dependent on the presence or absence of the lipid moiety. However, if Lipo-YcjN aggregation is primarily driven by the lipid moiety, we would expect ΔYcjN and Lipo-YcjN to elute at different volumes, indicative of lipid-dependent aggregation.

The elution profiles of recombinantly expressed ΔYcjN in the presence and absence of LMNG are shown in Figure 2B, right panel. In the presence of LMNG, ΔYcjN elutes at 85 mL (Figure 2B, dark green trace), while in its absence, ΔYcjN elutes at 84 mL (Figure 2B, light green trace). These peak elution volumes correspond well with the expected elution volume for a protein with a theoretical molecular weight of monomeric ΔYcjN (45,639 Da). Together, these results suggest that Lipo-YcjN, but not ΔYcjN, aggregates in solution. To confirm the aggregation of the Lipo-YcjN protein in solution, we performed DLS measurements on purified YcjN proteins and MBP as a control. The DLS measurements revealed that the hydrodynamic radius (Rh) of MBP was 3.0 *±* 0.1 nm (Figure S2.A), while the Rh of ΔYcjN was 3.3 *±* 0.1 nm (Figure S2.B). However, the Rh value of Lipo-YcjN was 70 *±* 5 nm, which is much larger than both MBP and ΔYcjN (Figure S2.C). Together, analyses of the DLS measurements and SEC profiles suggest that Lipo-YcjN aggregates in solution, possibly through hydrophobic lipid-lipid interactions.

In addition, a comparison of the YcjN SEC profiles reveals that the elution volumes of the major peaks of Lipo-YcjN appear to be sensitive to the presence of LMNG (Figure 2B, light and dark blue stars). In contrast, the elution volumes of ΔYcjN remain relatively similar with and without LMNG (Figure 2B, green traces). These data suggest that LMNG interacts with Lipo-YcjN aggregates, possibly solubilizing them into smaller Lipo-YcjN:LMNG micelle-like structures with slightly higher peak elution volumes (the difference between the major peaks indicated by star markers is 13 mL).

### Overall structure of **Δ**YcjN

To investigate the structural details of YcjN, we used X-ray crystallography to determine its three-dimensional structure. We chose the ΔYcjN protein for these crystallization studies because it does not aggregate in solution. We determined the first crystal structure of ΔYcjN at 1.95 Å resolution using the structure of an uncharacterized SBP from *Enterobacter cloacae* (PDB ID: 7V09) for molecular replacement. Data collection and structure refinement statistics are listed in Table 1. The crystal structure was solved with P1 symmetry and the asymmetric unit contained two molecules. A comparison of the two molecules in the asymmetric unit shows that they are very similar (RMSD of 0.182 Å across all 401 pairs).

**Table 1:** Data collection and refinement statistics.

| PDB ID | apo YcjN (8VQK) |
| --- | --- |
| **Data collection <sup>a</sup>** |  |
| Wavelength (Å) | 0.97946 |
| Space group | P 1 |
| Cell dimensions |  |
| a,b,c (Å) | 63.22 63.29 74.31 |
| $\alpha, \beta, \gamma$ (°) | 110.0 91.1 118.7 |
| Unique reflections | 61761 (6203) |
| Multiplicity | 3.9 |
| Completeness (%) | 91.19 (90.35) |
| Mean I/sigma(I) | 4.9 |
| CC <sub>1/2</sub> % | 99.8 (29.3) |
| **Refinement** |  |
| Resolution (Å) | 34.04 - 1.95 |
| R-work/ R-free | 0.22/ 0.26 |
| Number of non-hydrogen atoms | 6501 |
| macromolecules | 6139 |
| ligands | 43 |
| solvent | 319 |
| Protein residues | 803 |
| RMS(bonds) | 0.007 |
| RMS(angles) | 0.81 |
| Ramachandran |  |
| favored (%) | 98.25 |
| allowed (%) | 1.63 |
| outliers (%) | 0.13 |
| Rotamer outliers (%) | 2.23 |
| Clashscore | 5.91 |
| Average B-factor | 39.89 |

Numbers in parentheses correspond to the highest resolution shell

The ΔYcjN structure is composed of two *α*/*β* domains: the N-terminal domain (NTD) and the C-terminal domain (CTD) that can be divided into two subdomains (CTD1 and CTD2) (Figure 3, shown in teal, gray, and light blue, respectively). Each domain comprises two non-consecutive amino acid segments (Figure 3B). The NTD possesses a three-stranded *β*-sheet flanked by 11 *α*-helices (excluding linker segments). The CTD is composed of two subdomains (CTD1, residues 145-298 and residues 359-383; CTD2, residues 384-430) and is composed of three *β*-strands and 13 *α*-helices. The NTD and CTD are connected by three segments, including *β*7, *β*4, and *α*15. An analysis of the binding pocket revealed that there is no discernible electron density corresponding to a bound ligand deep within the binding pocket. However, a smaller electron density was detected at the edge of the binding pocket near tyrosine 95. We assign this electron density to polyethylene glycol (PEG) since it was included as an additive in the crystallization experiment (Figure S3.A).

**Figure 3:**
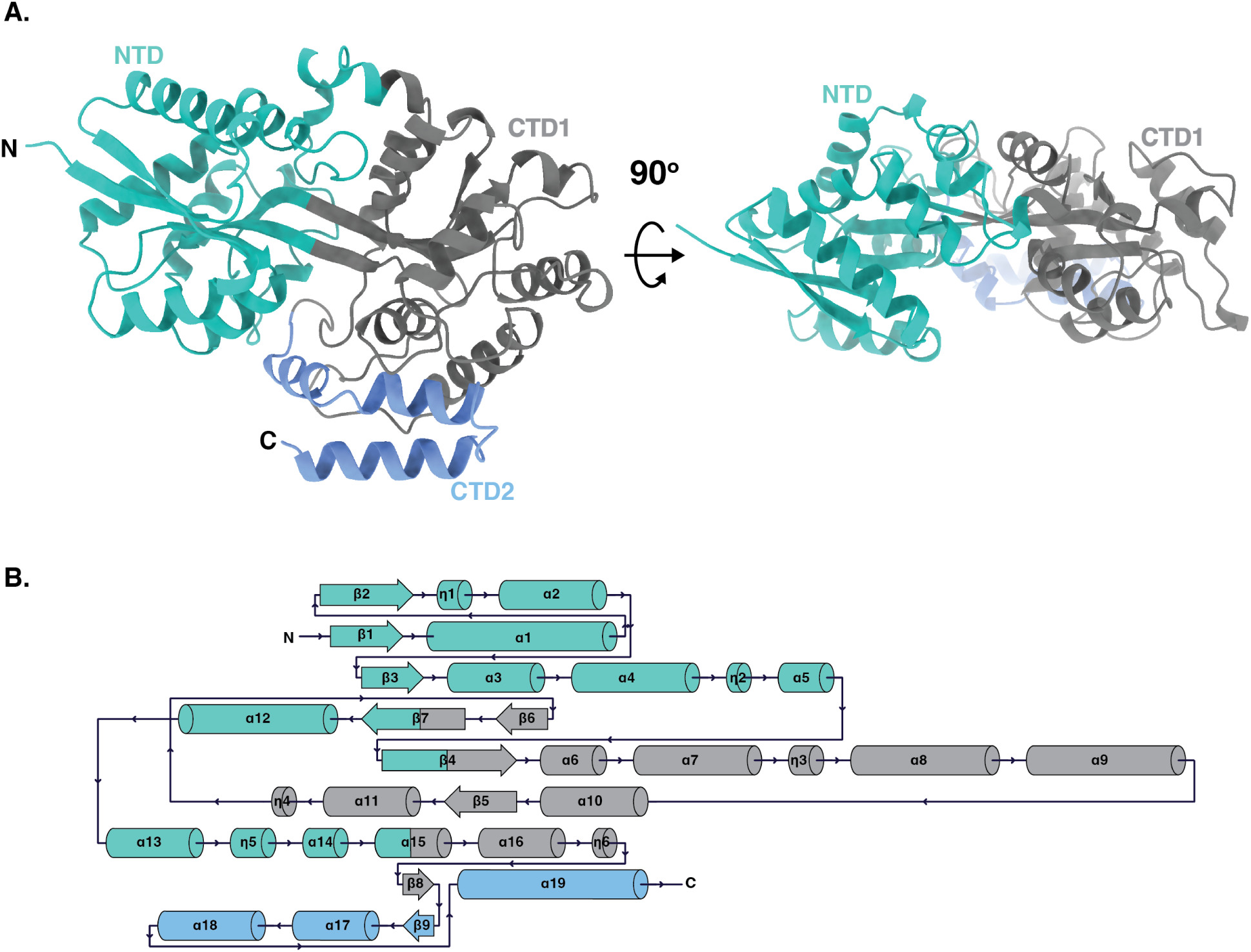
Overall structure of *E. coli* YcjN. A. Ribbon representation of the YcjN structure. The same structure is shown with a 90*^◦^*C x-axis rotation. B. Schematic of secondary structure. Subdomains NTD, CTD1, and CTD2 are colored teal, gray, and light blue, respectively.

### Comparison of YcjN to other SBPs

A structural similarity search using the Dali server^14^ revealed that YcjN is most closely related to putative carbohydrate-binding proteins. Top results from the search include various sugar binding proteins (Figure 4), such as tmMBP3^15^ [Z-score, 41.6 ; RMSD 2.4 Å for 393 residues, 22% identity], mtUgpB^16^ [Z-score, 38.7; RMSD 2.3 Å for 403 residues, 19% identity], xaMalE^17^ [Z-score, 38.6; RMSD 2.6 Å for 397 residues, 18% identity], TTHA0356^18^ [Z-score, 38.5; RMSD 2.5 *Å* for 416 residues, 14% identity], and sgGacH^19^ [Z-score, 38.4; RMSD 2.5 Å for 391 residues, 19% identity]. Notably, all of these proteins contain a distinct CTD2 domain (Figure 4A, blue helices) and have molecular weights greater than 40 kDa (Figure 4D). These characteristics are common among proteins belonging to subcluster D-I, which includes MBP.^20,21^ SBPs that bind carbohydrates can also be members of subcluster B-I.^11^ However, these proteins lack the CTD2 domain, making them structurally distinct from subcluster D-I proteins (Figure 4C). Due to YcjN’s molecular weight and the presence of the CTD2 domain, we group YcjN into subcluster D-I.

**Figure 4:**
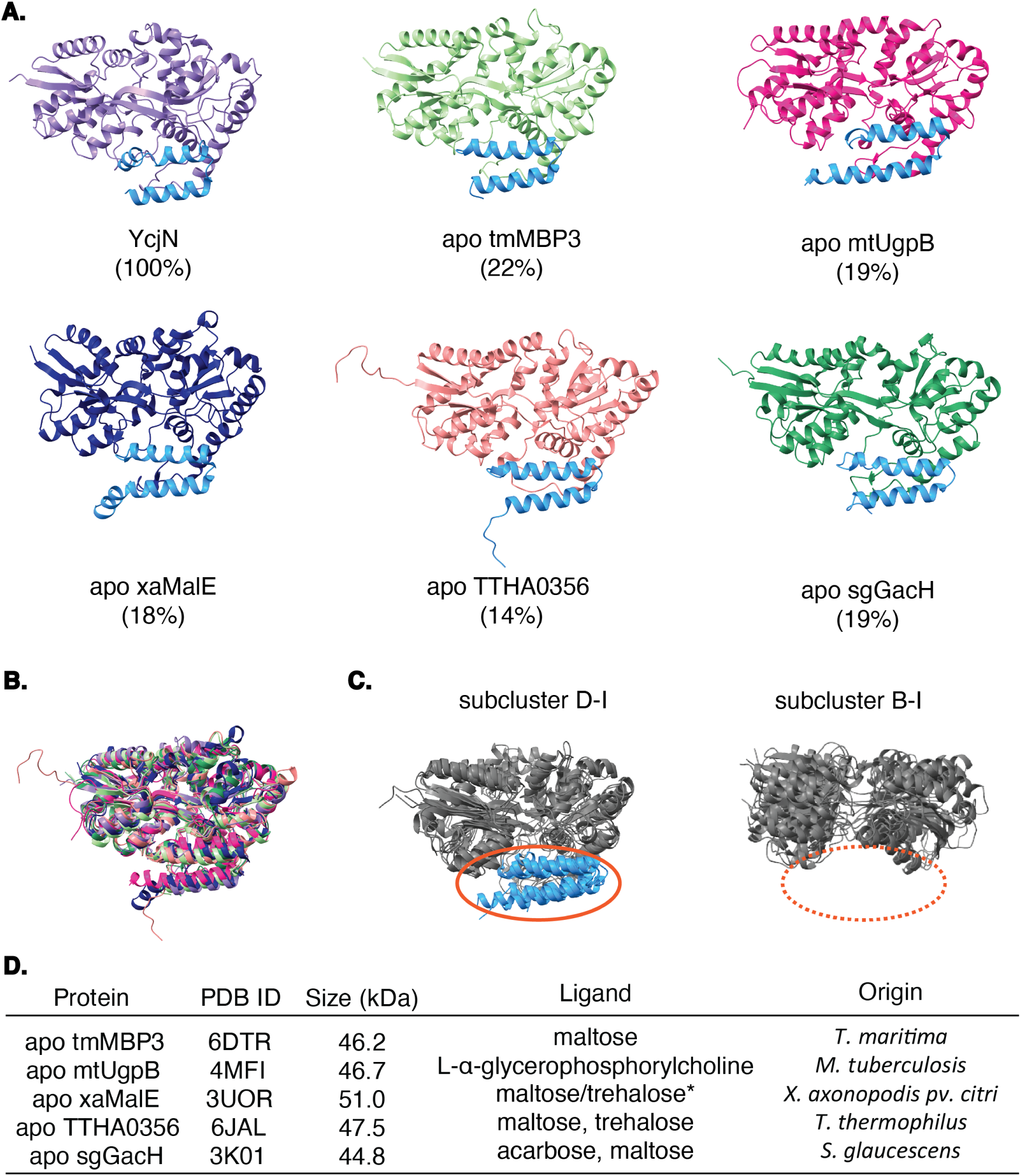
Comparison of YcjN to other SBPs identified using the Dali Server. A. Structures and percent identities to YcjN. The CTD2 domain of each protein is colored blue. B. Superposition of SBPs. Each protein color corresponds to that of the protein in panel A. C. SBP subcluster categorization of the aligned proteins. All CTD2 domains (blue) present in subcluster D-I (1ANF, 2ZYK, 4AQ4, and 2UVI) are absent in proteins within subcluster B-I (2FW0, 1GCG, 1URP, 1GUB) that also bind carbohydrates. D. Table of proteins, their size, ligand (*putative), and species of origin.

Top hits from the Dali search also have a low percent identity to YcjN (*<* 25%) (Figure 4A) and the volumes of their binding pockets vary (551 Å^3^ to 1432 Å^3^, determined using CASTp 3.0 with a probe sphere 1.4 Å) (Figure S3.B). Furthermore, a sequence comparison did not reveal any strict identity of amino acid residues within the binding pockets for these proteins (Figure S4). Despite variations in binding pocket volumes and low percent identities, three of the five proteins structurally similar to YcjN bind carbohydrates, including maltose, while another protein binds glycerophosphorylcholine (Figure 4D). Therefore, it is plausible that YcjN could also bind similar ligands. However, experimental validation is necessary to identify YcjN’s ligand.

### NanoDSF screening to identify YcjN ligands

Unlike some SBPs, ΔYcjN did not crystallize bound to a ligand deep within its binding pocket. Therefore, to determine the ligand binding profile of ΔYcjN, we employed a thermal shift assay using nano-Differential Scanning Fluorimetry (nanoDSF). This method uses tryptophan/tyrosine fluorescence at 330 nm and 350 nm to monitor protein thermal unfolding. These profiles can then be used to determine the protein’s denaturation midpoint (Tm), which reflects the stability of the protein or protein-ligand complex. In our experiments, the protein’s Tm was measured in the presence and absence of potential ligands and the ΔTm was calculated. The molecule was then classified as a ligand if the calculated ΔTm was greater than 2*^◦^*C.

Since YcjN is part of a gene cluster encoding 12 proteins involved in carbohydrate metabolism, we included a range of commercially available monosaccharides and polysaccharides in our screen (Figure 5). Considering the structural similarity of ΔYcjN to tmMBP3 and mtUgpB, we also included maltose and glycerophosphorylcholine in our screen. Additionally, this screen included kojibiose because it is a known substrate for YcjT.^12^ As a positive and negative control, we calculated the ΔTm of MBP in the presence of two known ligands (maltose and maltotriose) and two non-ligands (lactose and sucrose), respectively.^22^ As expected, the ΔTm of MBP increased by 8.6*^◦^*C and 9.1*^◦^*C in the presence of maltose and maltotriose, respectively, while in the presence of non-ligands, lactose and sucrose, the ΔTm increased by less than 0.5*^◦^*C. Next, we monitored the thermal stability of ΔYcjN in the presence and absence of various potential ligands. These ligands did not produce a detectable thermostabilizing effect on ΔYcjN, with the ΔTm remaining below 0.5*^◦^*C for all screened ligands. Together, these data suggest that ΔYcjN’s ligands may not have been part of our screen or had no detectable thermostabilizing effects on ΔYcjN. As a result, the identities of ΔYcjN’s ligands remains unknown.

**Figure 5:**
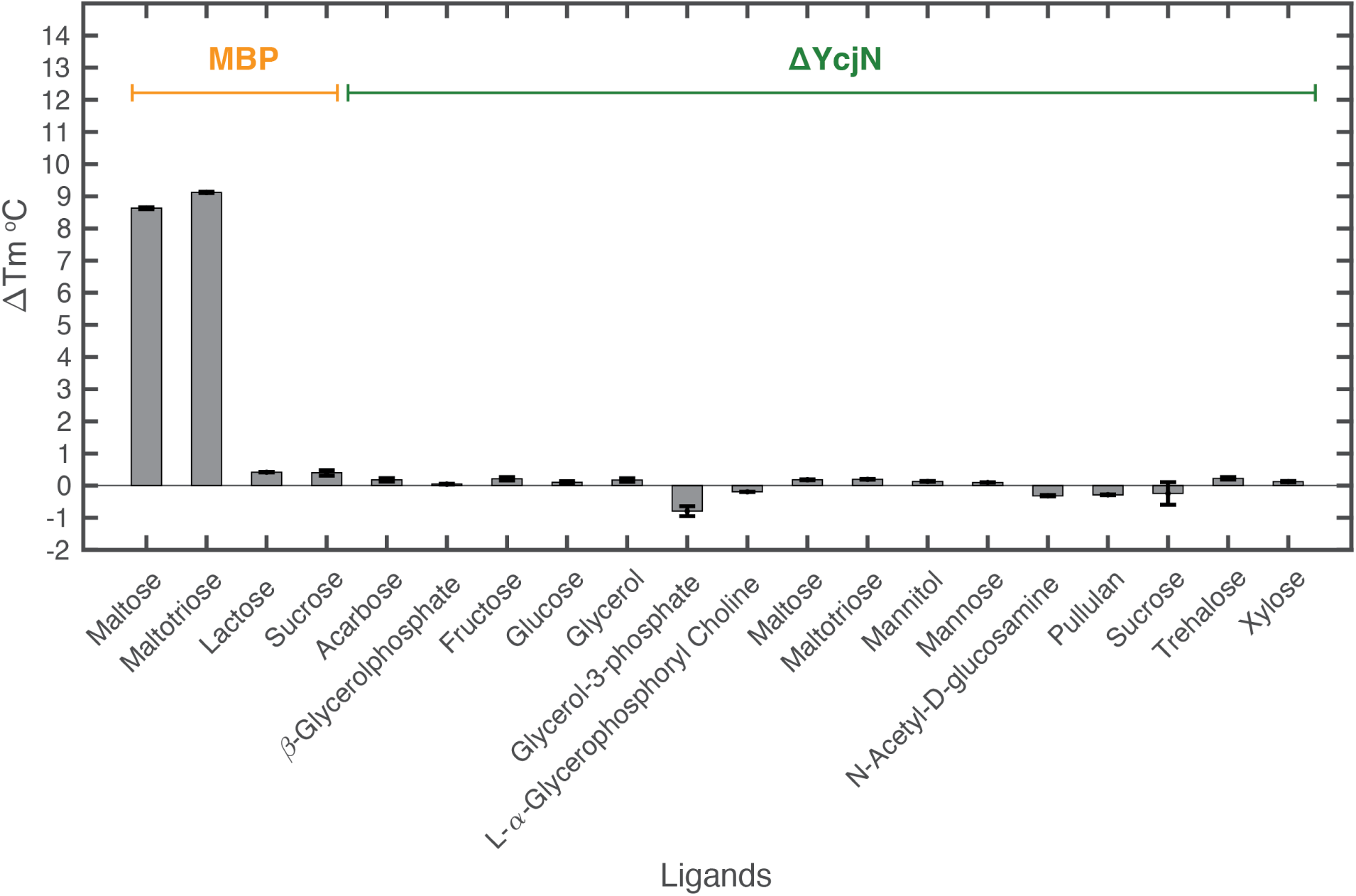
NanoDSF thermal shift assay for MBP and ΔYcjN. Bar graph displaying ΔTm for MBP and YcjN with potential ligands. All measurements were performed using final protein concentrations of 1 mg/ml and ligand concentrations of 5 mM. Values represent the calculated mean *±* standard error of three technical replicates.

### Bacteria and Archaea contain non-lipidated and lipidated YcjN-like proteins

To determine whether lipidated YcjN-like proteins are exclusively present in *Enterobacteriaceae* such as *E. coli*, we investigated the taxonomic distribution and putative lipidation state of these proteins using bioinformatic tools. Specifically, to identify YcjN homologs, we used the Enzyme Function Initiative - Enzyme Similarity Tool (EFI-EST) program to create a Sequence Similarity Network (SSN) for YcjN and its closest homologs in the UniRef90 database. In the SSN, each protein is depicted as a node (circles), and if they exhibit pairwise protein sequence similarity, the nodes are connected by edges (lines). To identify putative lipidated and non-lipidated forms of YcjN homologs, we used SignalP 6.0. This algorithm predicts which enzymes are likely to be involved in the posttranslational modification of immature proteins based on the protein’s amino acid sequence. Nodes in the SSN were manually annotated and colored as lipoproteins in purple, non-lipidated proteins in light blue, and other in orange, if SignalP 6.0 predicted that they were processed by SPase I, SPase II, or classified as other, respectively. This analysis also produced a sunburst diagram with the taxonomic distribution of YcjN-like proteins (Figure 6A). An inspection of this diagram suggests that YcjN homologs are primarily found in both Archaea and Bacteria. In addition, inspection of the SSN reveals that YcjN could be in a lipidated or non-lipidated form, suggesting that lipidation may not be essential for YcjN’s function. Furthermore, these proteins are separated into several distinct clusters, with either lipidated or non-lipidated YcjN protein forms dominating each cluster (Figure 6B,C). For example, YcjN from *E. coli* (Figure 6B, cluster 3, star marker) is predicted to share a high sequence similarity to other putative YcjN lipoproteins.

**Figure 6:**
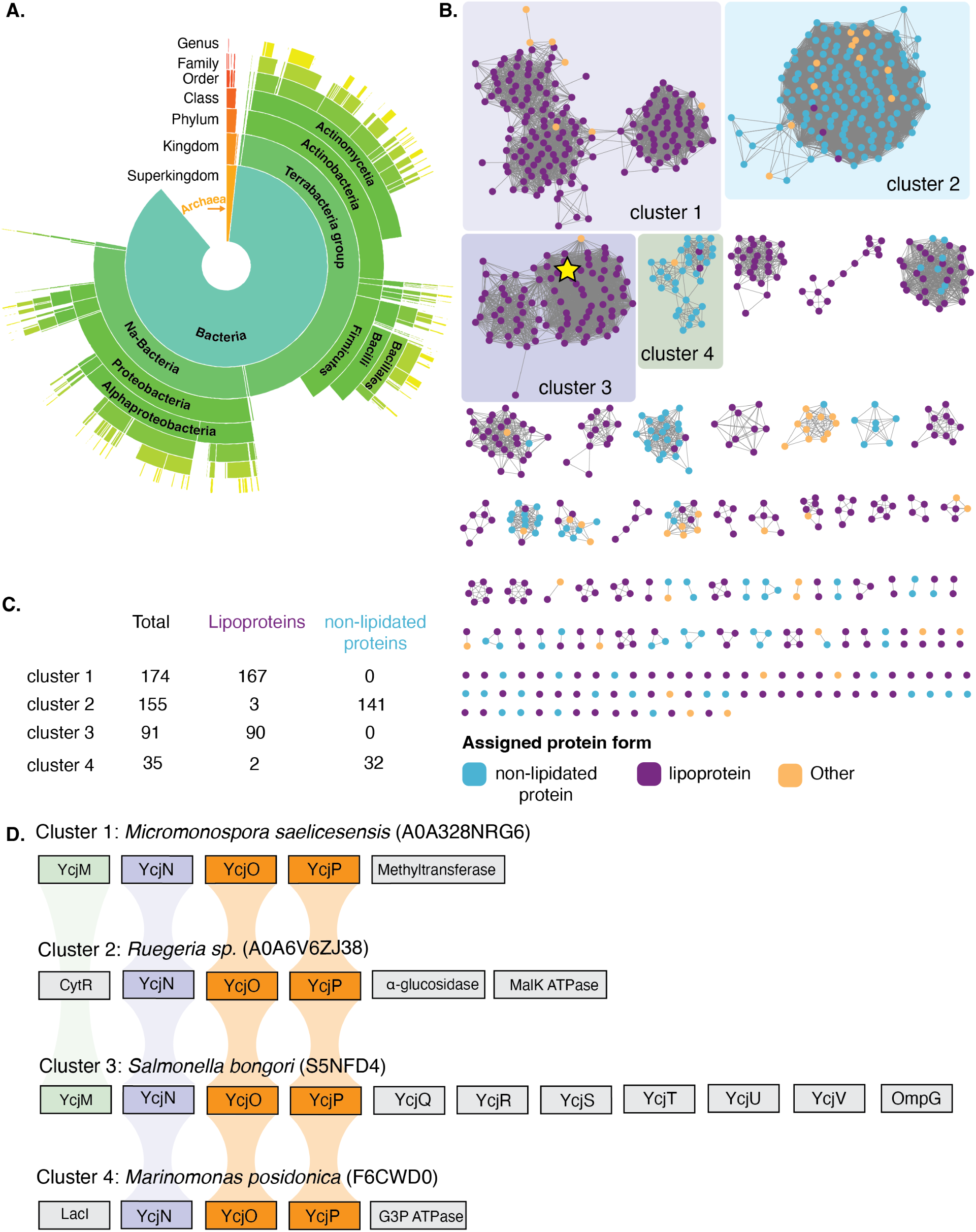
Bioinformatic analysis of YcjN. A. Sunburst diagram of the taxonomic distribution of YcjN homologs. B. The SSN diagram. C. Total number of proteins labeled as lipoproteins and non-lipidated proteins in clusters 1-4. D. Representative diagrams of the putative proteins within clusters 1-4 that are frequently encoded near *YcjN* as determined using the EFI-GNT tool. Putative YcjM, YcjN, YcjO/YcjP proteins are colored in green, purple, and orange, respectively.

Next, we analyzed the genomic context of four clusters (1-4) to investigate the co-occurrence of genes near *YcjN*. This analysis showed that cluster 3 exhibited the most similar co-occurrence of genes to that of *E. coli*, with genes encoding proteins YcjM, YcjN, YcjO, YcjP, YcjQ, YcjR, YcjS, YcjT, YcjU, YcjV, and OmpG (Figure 6D). This observation is not surprising since *E. coli* YcjN belongs to cluster 3 and shares the highest protein similarity to YcjN proteins within this group. Moreover, all four clusters contained putative genes encoding proteins YcjO and YcjP (Figure 6D, orange), and two of the four clusters also included a gene that encodes the YcjM protein (Figure 6D, green). These data suggest that the gene encoding YcjN often co-occurs with genes encoding the transmembrane domains (TMDs) of an ABC transporter and, occasionally, next to a gene encoding a sugar phosphorylase. A closer inspection of cluster 3 reveals that it comprises various bacteria from the phylum *Proteobacteria* such as *Citrobacter farmeri*, *Shigella boydii*, and *Enterobacter cloacae*, as well as from the *Firmicutes* phylum, including *Enterococcus pseudoavium*, *Listeria fleischmannii*, and *Bacillus sp*. Many of these bacteria have been identified in food, soil, and humans/animals. Notably, none of the proteins within this cluster have been experimentally characterized and their ligands remain unknown. Together, these data suggest that YcjN may be involved in carbohydrate import with the closest homologs of YcjN found in several bacteria that inhabit animals or soil.

## Discussion

Nutrient acquisition is a fundamental process essential for bacterial growth and survival. To scavenge nutrients, bacteria employ several mechanisms, including the active transport of molecules using SBPs and their cognate ABC transporters. *E. coli* possesses several SBPs with distinct ligand binding profiles, allowing the bacterium to adapt to environments with varying nutrient compositions. To gain structural insight into the SBP YcjN, we recombinantly overexpressed, purified, and characterized lipidated and non-lipidated YcjN from *E. coli*. Specifically, we report the first crystal structure of ΔYcjN determined at 1.95 Å resolution. In the crystal structure, the binding pocket is empty except for a PEG molecule bound to the outer edge of the binding pocket. Most likely, this PEG molecule was introduced during the crystallization experiment. A structural similarity search showed that YcjN closely resembles sugar-binding proteins and an analysis of its structure revealed that it can be classified into subcluster D-I, which comprises SBPs with molecular weights greater than 40 kDa and a distinct CTD2 domain. Notably, this group includes MBP.

Unlike some SBPs, ΔYcjN did not crystallize bound to a ligand deep within its binding pocket. Therefore, to determine the ligand binding profile of ΔYcjN, we employed a nanoDSF thermal shift ligand binding assay. Despite our efforts, our screen did not reveal any ligand that produced a thermostabilizing effect greater than 2 *^◦^*C. Specifically, no binding was detected for known ligands of structurally similar SBPs, such as maltose and glycerophosphorylcholine, as well as kojibiose, a substrate for YcjT. Importantly, the lack of a thermostabilizing effect does not indicate the absence of ligand binding. Because our assay does not measure ligand binding directly, it’s possible that YcjN binds these ligands but does not produce a measurable thermostabilizing effect. However, we believe that it is more likely that YcjN’s endogenous ligand was absent from our screen because YcjN shares structural similarity to MBP which, in our assay, clearly showed a thermostabilizing effect in the presence of known ligands maltose and maltotriose. Given that our bioinformatic analysis suggests that YcjN-like proteins are found in food, animals, or soil dwelling bacteria with its gene co-occurring proximal to a sugar phosphorylase, we speculate that its ligand could be a plant-derived carbohydrate, glycolipid, or metabolite. Consequently, future efforts should consider employing an alternative screening method or broadening the classes of ligands in the screen.

Our results also show that Lipo-YcjN aggregates in solution, whereas its non-lipidated form does not. Similar observations have been made for other lipoproteins. For example, the SEC elution profiles of lipidated *N. meningitidis* MetQ and lipidated *B. burgdorferi* OspA reveal lower peak elution volumes compared to their non-lipidated counterparts.^5,23^ Moreover, studies on the lipoprotein *N. meningitidis* fHbp using size-exclusion chromatography with multi-angle light scattering revealed a molecular weight of 660 kDa, which is much higher than the theoretical weight of its monomer (28 kDa).^24^ Together, these findings suggest that lipid-mediated aggregation may be an intrinsic property of bacterial lipoproteins. While the biological relevance of lipoprotein aggregation within the cell may be limited since lipoproteins anchor to cellular membranes in bacteria, understanding lipoprotein properties in solution may be important for in vitro studies. For example, *B. burgdorferi* OspA and *N. meningitidis* fHbp are two recombinantly expressed lipoprotein antigens included in vaccine formulations. Importantly, their lipid moieties play a key role in inducing a robust immune response. ^23,25^ Therefore, a comprehensive understanding of lipoprotein aggregation in solution may be important for the effective separation of lipidated and non-lipidated protein forms using chromatography techniques, a key step in isolating lipidated antigens for vaccine development.

Our study also details protocols for purifying Lipo-YcjN both with and without LMNG. Previous studies have used a variety of detergents, including Triton X-100 and DDM, to purify histidine-tagged lipoproteins via affinity chromatography.^5,26^ However, here we demonstrate that Lipo-YcjN can be purified in the absence of detergent. This result suggests that detergents are not an essential component for the purification of all lipoproteins. In addition, analysis of the SEC profiles for Lipo-YcjN in the presence and absence of LMNG revealed a higher peak elution volume compared to that of its non-lipidated form. This data suggests that LMNG may solubilize lipoprotein aggregates, possibly into smaller lipoprotein-LMNG mixed micelles. However, the interactions between detergents and lipoproteins remain poorly understood and require further investigation.

Another key finding of our study is that recombinantly produced YcjN is predominantly observed in its diacylated form rather than in its triacylated form. This outcome is surprising, because *E. coli* contains all three enzymes to triacylate lipoproteins (Lgt, SPaseII and Lnt), and our previous work on the recombinant overexpression of *N. meningitidis* MetQ, which used the same expression protocols, detected only triacylated *N. meningitidis* MetQ. Other researchers have found that expression conditions, such as media and pH, can influence the production of diacylated versus triacylated lipoprotein forms.^27^ Therefore, our results, in light of previous studies, raise several questions about the biosynthesis and processing of lipoproteins: Could the expression of specific proteins influence environmental conditions, thereby favoring the formation of diacylated lipoproteins? Alternatively, could the N-terminal amino acids, such as the signal peptide, trigger an early exit from the lipoprotein maturation pathway? Are diacylated lipoproteins an uncharacterized form in *E. coli*, or are they just artifacts of recombinant overexpression methods? Given that these possibilities are not mutually exclusive, two or more factors could help explain the preference for endogenous and recombinant production of diacylated over triacylated lipoproteins in *E. coli*.

In summary, this study contributes to our overall understanding of Lipo-YcjN, providing a foundation for future research on bacterial lipoproteins. These findings are particularly relevant for future studies that aim to characterize bacterial lipoproteins in solution and to help advance methods for the production of recombinant lipidated proteins for vaccine formulations.

## Experimental Procedures

### Cloning, expression, and purification of YcjN and MBP proteins

The amino acid sequence of YcjN was obtained from *E. coli* K-12 (gene ID 945696; UniProt ID P76042). To produce YcjN constructs, the DNA sequence encoding YcjN was inserted into a pET21b(+) ampicillin-resistant vector between NdeI and XhoI sites and under the control of a T7 promoter (GenScript, USA). Two constructs were created: SP-YcjN-10H, 6H-ΔYcjN (residues 1-21 removed). To aid in purification, nucleotide sequences encoding either decahistidine (10H) or hexahistidine (6H) were added to the N- or C-terminus of the *YcjN* genes. Similarly, the amino acid sequence of MBP was obtained from *E. coli* K-12 (gene ID 948538; UniProt ID P0AEX9). In this construct, residues 1-26 encoding the SP were replaced with a 6H. All proteins were expressed in *E. coli* BL21 (DE3) gold cells (Agilent, USA) using ZYM-5052 autoinduction media^28^ containing 100 mg/L ampicillin at 37 *^◦^*C for 30 h. Cells were harvested by centrifugation at 4785 xg (JLA 8.1; 5000 rpm; Beckman Coulter, USA) for 15 min and the cell paste was flash frozen in liquid nitrogen for storage at -80 *^◦^*C.

To purify YcjN and MBP proteins, *∼*10 g of cell paste was thawed at room temperature (RT), then placed in solution of 100 mL of 25 mM Tris HCl pH 7.5 and 100 mM NaCl, 40 mg of lysozyme, 4 mg of DNase, and one cOmplete protease inhibitor cocktail tablet (Sigma-Aldrich, USA). The suspended cells were disrupted using a Microfluidizer (Microfluidics, USA) and cell debris was removed by centrifugation at 7823 xg (Ti45; 10,000 rpm; Beckman Coulter, USA) for 30 min. Imidazole was then added to the lysate to remove nonspecific protein binders during affinity purification (25mM imidazole for 6H constructs; 70 mM imidazole for 10H constructs). Proteins were purified using a 5 mL HisTrap HP column (Cytivia, USA) pre-equilibrated with 25 mM Tris HCl pH 7.5 and 100 mM NaCl and eluted with a solution of 25 mM Tris HCl pH 7.5, 100 mM NaCl, and 300 mM Imidazole. The eluted proteins were then loaded onto a Hiload 16/600 Superdex 200 pg (Cytiva, USA) pre-equilibrated with 25 mM Tris HCl pH 7.5 and 100 mM NaCl. YcjN proteins purified in the presence of LMNG were obtained using the same protocol with the following modifications: After cell disruption, LMNG was added to the lysate at a final concentration of 1% and incubated at 4 *^◦^*C for 3 h. Cell debris was removed by centrifugation at 94,834 xg (Ti45; 35,000 rpm; Beckman Coulter, USA). For affinity and size exclusion chromatarography, the same protocols were used with buffers containing LMNG added to a final concentration of 0.01%. For all proteins, the peak fractions were combined, frozen in liquid nitrogen, and stored at -80 *^◦^*C until thawed. Mass analysis of intact YcjN proteins was performed as previously described.^5^ The mass spectrum graph was imported into Adobe Illustrator (CC 2024), where text and font size were modified to increase readability. Graphs were generated in Matlab (version R2022b).

### Dynamic light scattering

Dynamic light scattering measurements of YcjN and MBP proteins were performed using SEC peak fractions using the default Measure Size option on a DynaPro Nanostar II (Wyatt Technology Corporation, USA). A total of five measurements per sample were acquired, then DLS profiles and hydrodynamic radii values were exported to Matlab. This program was then used to calculate the mean and standard deviation of the measurements and to plot the representative profiles.

### Crystallization, data collection, structural determination, and structural analysis of **Δ**YcjN

The pooled factions of ΔYcjN were concentrated to 175 mg/mL for crystallization trials. The ΔYcjN protein was crystallized at 16*^◦^*C by the sitting-drop vapor diffusion method using a precipitant solution from the BCS Screen (Molecular Dimensions Inc., USA) consisting of 0.01 M CdCl_2_, 0.2 M NH_4_SO_4_, 0.1 M HEPES pH 7.5, and 25 % w/v mix consisting of PEG 3350, PEG 4000, PEG 2000, and PEG 5000 MME. The crystals were cryocooled in liquid nitrogen after dipping them into a cryoprotectant solution composed by the precipitant solution supplemented with 25 % PEG 400. The X-ray diffraction data was collected on SSRL beamline 12-1 at the Stanford Synchrotron Radiation Lightsource (SSRL) at the SLAC National Accelerator Laboratory, processed with the XDS package,^29^ and scaled and merged with SCALA.^30^ The ΔYcjN protein crystal belonged to the P1 space group and the structure was determined by molecular replacement (MR) with PHASER^31^ using the coordinates of *Enterobacter cloacae* (PDB ID: 7V09, 75 % identity) an uncharacterized protein as the search model. The model was manually adjusted using Coot^32^ and refined using Phenix.refine.^33^ The atomic coordinates and structure factors have been deposited in the RCSB Protein Data Bank (8VQK). Figures were created by employing PDBsum^34^ to generate secondary structural schematics and ESPript 3.0^35^ to obtain secondary structure labels. These elements were manually integrated using Adobe Illustrator to produce the final figure.

The visualization of the protein binding pockets and the identification of the residues lining the pockets were performed by the CASTp 3.0 server, using default values. ^36^ Multiple sequence alignment was performed using the align functionality in UniProt and ESPript 3.0 was used to visualize the alignment in the Black and White scheme. In the final sequence alignment, the residues lining the pockets were manually colored in red using Adobe Illustrator.

Visualization of YcjN electron density maps was performed using ChimeraX^37^ or the Moorhen website (https://moorhen.org). The protein similarity search was performed using the DALI server against the PDB90 database.^14^ The top protein hit, 7V09 with 75 % identity, was excluded from the structural comparison since it was used as the model for molecular replacement.

### NanoDSF

To remove potential endogenously bound ligands, we purified ΔYcjN and MBP and subjected both proteins to dialysis, using a protocol similar to that previously described. ^38^ The protein ΔYcjN was chosen for these studies because the thermal profile of lipid-modified YcjN may be a convolution of aggregate disassembly and protein unfolding, which could pose challenge in data analysis. Briefly, after purification, dialysis was performed using a 12 mL Slide-A-Lyzer Dialysis Cassette with a molecular weight cut-off of 3.5K (ThermoFisher Scientific, USA). The dialysis was conducted at 4 *^◦^*C for a total duration of 48 h. During this period, proteins were dialyzed against 2 L of a solution containing 25 mM Tris HCl at pH 7.5 and 100 mM NaCl, and the dialysis solution was replaced once after 24 h.

NanoDSF was performed on a Prometheus Panta (NanoTemper Technologies Inc., USA). The 330/350 nm fluorescence ratio was recorded between 25 *^◦^*C and 70 *^◦^*C at 1 *^◦^*C/min. Measurements were made using Prometheus nanoDSF Grade standard capillaries using excitation power of 20 %. All measurements were performed using 2 mg/mL protein concentrations and 10 mM ligand stock solutions in 25 mM Tris HCl pH 7.5 and 100 mM NaCl. For ligand binding experiments, protein stock solutions were mixed with an equal volume of ligand stock solution at a 1:1 v/v ratio to give a final solution of 1 mg/mL with 5 mM ligand in 25 mM Tris HCl pH 7.5, 100 mM NaCl. The ligand free sample was prepared using 1:1 v/v ratio with buffer to give a final solution of 1 mg/mL protein concentration. All sugars, glycerol, *β*-glycerophosphate disodium salt hydrate, *sn*-glycerol-3-phosphate bis(cyclohexylammonium) salt, and *sn*-glycero-3-phosphocholine 1:1 cadmium chloride adduct were all purchased from Sigma-Aldrich, with the exception of acarbose, which was purchased from Thermo Fisher Scientific.

### Bioinformatics and data analysis

The sequences were obtained using EFI-EST^39,40^ using the BLAST retrieval option (Option A) with the Uniref90 database and the FASTA sequence of the YcjN protein, which was obtained from UniProt.^41^ Taxonomy categories included the superkingdoms of Bacteria, Eukaryota, Viruses, and Archaea. The metagenome species and the phylum that fell in the unclassified fungi category were excluded. The final SSN was created using an alignment score threshold of 150 and a sequence restriction minimum of 400. The SSN contained 909 nodes with more than 40 % sequence similarity to the YcjN protein. SignalP 6.0 was then used to analyze the FASTA sequences of the SSN.^42^ SignalP 6.0 produced a summary of 896 predicted sequences, which included their respective type of signal peptide. Proteins predicted to be processed with SPase I or SPase II, and other were annotated as non-lipidated proteins, lipoproteins, or other, respectively. Protein sequences that failed the SignalP 6.0 analysis were removed from the SSN. The final SSN contained 896 protein sequences (Supplementary Data 1) and was visualized using the organic layout on Cytoscape (https://cytoscape.org). The taxonomy distribution sunburst was imported into Adobe Illustrator, where text and font size were modified to increase readability. EFI-GNN tools were used to analyze the genome context of YcjN-like proteins. The output of the EFI-GNN analysis was used to manually create protein diagrams in Adobe Illustrator. This article contains supporting information.

## Acknowledgement

The authors thank Dr. Qianqiao Liu for proofreading and critical reading of the manuscript and Stanford University for funding and facility use. This work utilized the NanoTemper Prometheus Panta dynamic light scattering (DLS) and nanoDSF system that was purchased with funding from Stanford c-SHARP Program. Authors thank the staff at SSRL, SLAC National Accelerator Laboratory for beamline assistance. Use of the Stanford Synchrotron Radiation Lightsource, SLAC National Accelerator Laboratory, is supported by the U.S. Department of Energy, Office of Science, Office of Basic Energy Sciences under Contract No. DE-AC02-76SF00515. The SSRL Structural Molecular Biology Program is supported by the DOE Office of Biological and Environmental Research, and by the National Institutes of Health, National Institute of General Medical Sciences (P30GM133894). The contents of this publication are solely the responsibility of the authors and do not necessarily represent the official views of NIGMS or NIH.

## Abbreviations

(ABC): ATP-binding cassette
(TMD): transmembrane domain
(NBD): nucleotide binding domain
(SBP): substrate binding proteins
(IM): inner membrane
(OM): outer membrane
(SP): signal peptide
(SPase I): signal peptidase
(Lgt): phosphatidylglycerol:prolipoprotein diacylglyceryl transferase
(SPase II): Type II signal peptidase
(Lnt): N-acyltransferase
(*E. coli*): *Escherichia coli*
(LC-MS): liquid chromatography mass spectrometry
(DLS): dynamic light scattering
(MBP): maltose binding protein
(SEC): size-exclusion chromatography
(LMNG): detergent 2,2-didecylpropane-1,3-bis-*β*-D-maltopyranoside
(NTD): N-terminal domain
(CTD): C-terminal domain
(PEG): polyethylene glycol
(nanoDSF): nano-Differential Scanning Fluorimetry
(Tm): protein denaturation midpoint
(EFI-EST): Enzyme Function Initiative - Enzyme Similarity Tool
(SSN): Sequence Similarity Network

## Supporting Information Available

**Figure S1:**
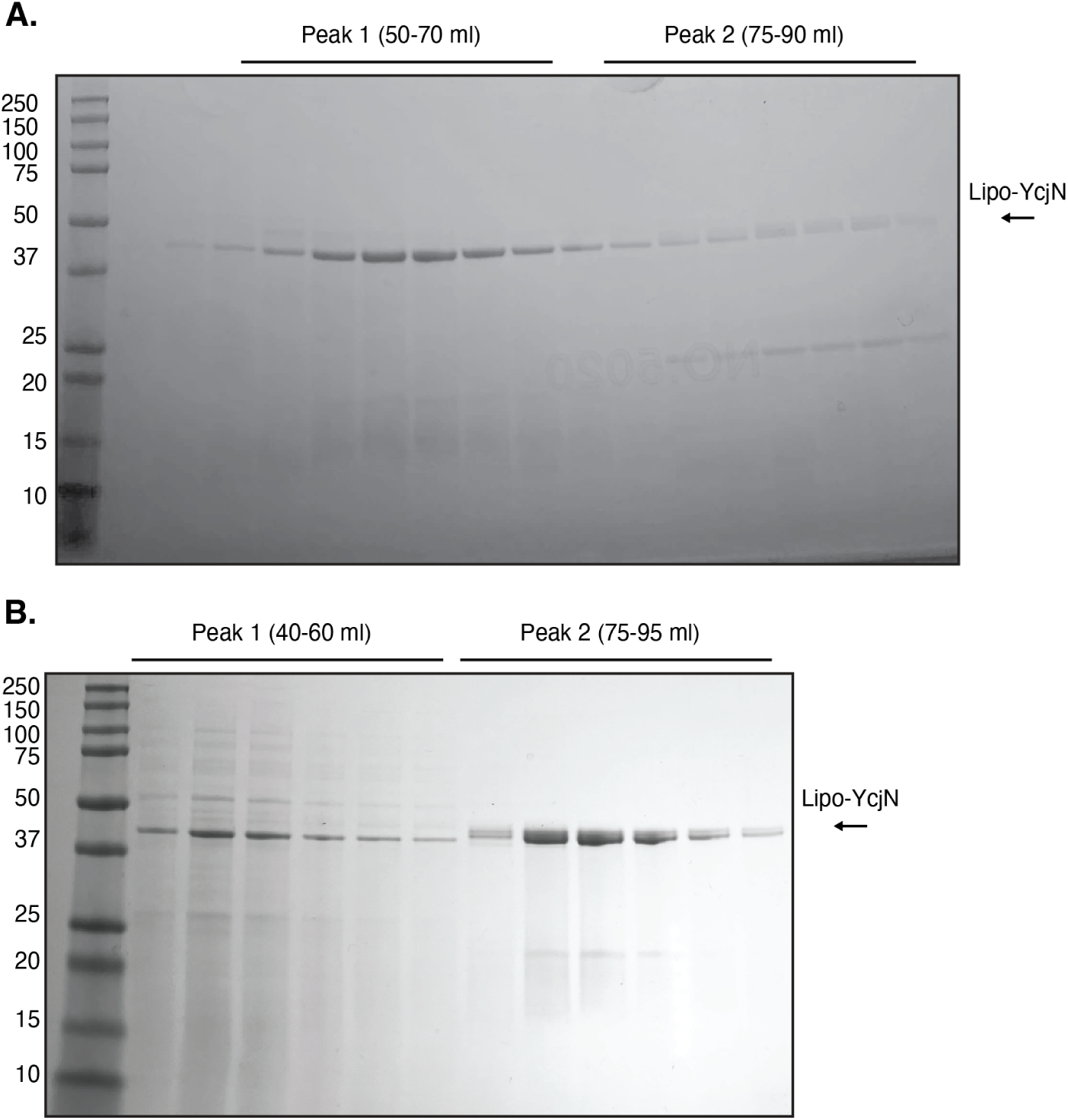
Analysis of SEC peak fractions using SDS-PAGE with ReadyBlue protein gel stain. A. Lipo-YcjN purified in the presence and B. absence of LMNG.

**Figure S2:**
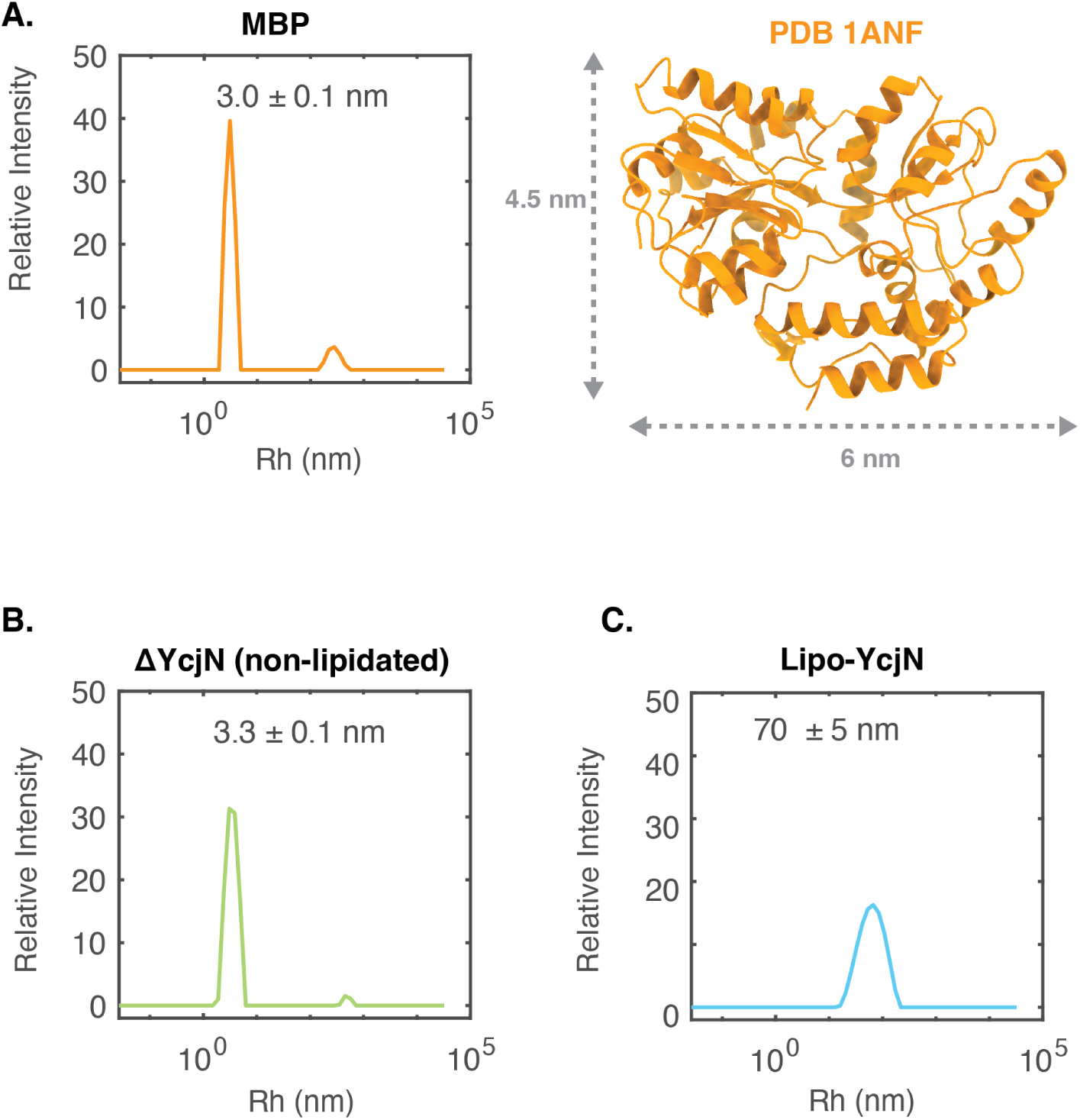
DLS analysis of YcjN and MBP proteins. A. DLS plot of MBP (left panel) and structure of MBP shown in ribbon representation (right panel). Approximate measurements of height and width are indicated by grey arrows. B. DLS plots for ΔYcjN and C. Lipo-YcjN purified in the absence of detergent (n=3 individual measurements, values shown as mean *±* SD).

**Figure S3:**
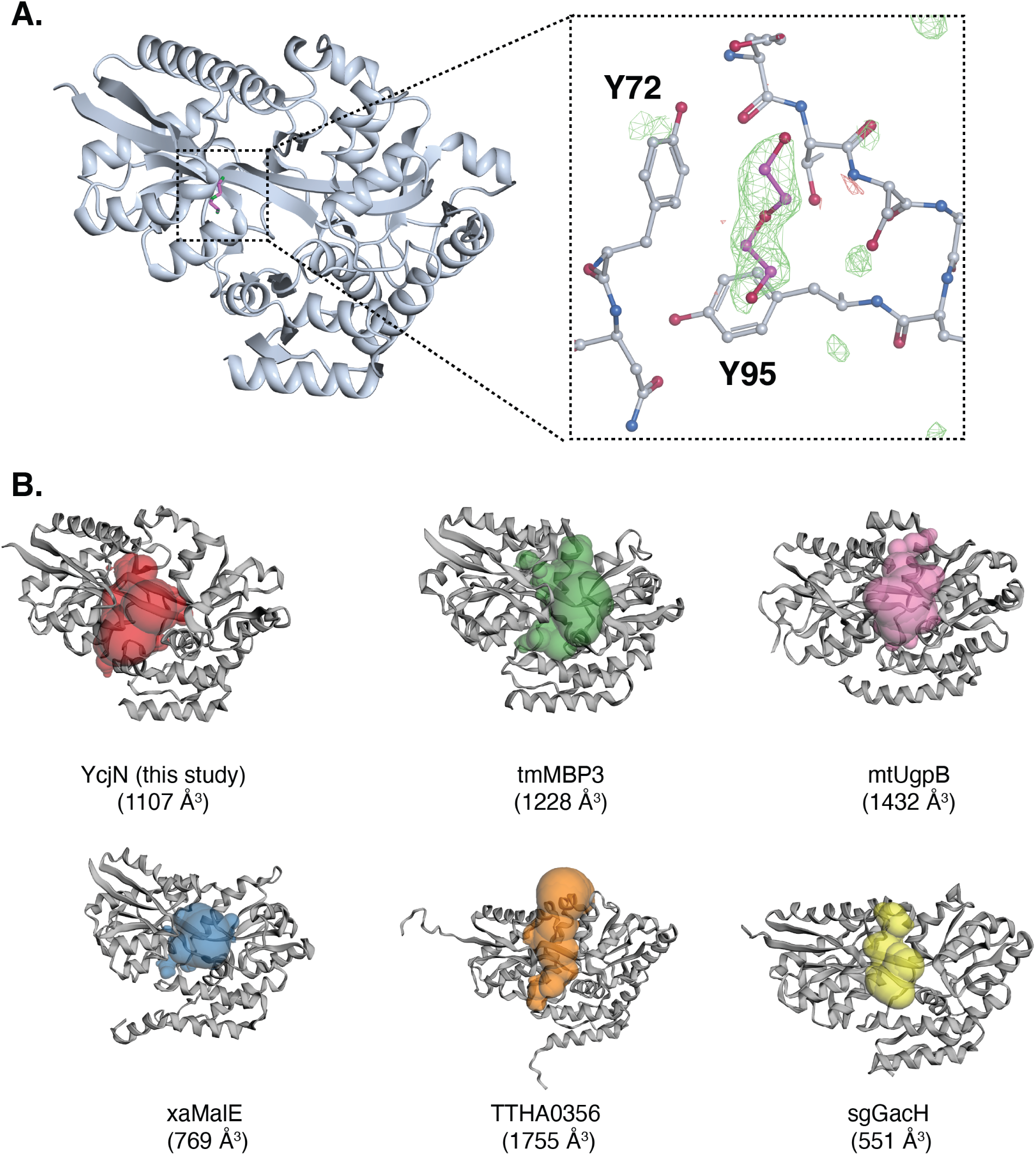
Ligand binding pockets of YcjN and YcjN homologs. A. Overall structure of YcjN (left panel) and an enlarged view of the binding pocket (right panel). In the enlarged view, the molecular model is depicted in stick representation. The Fo-Fc (2.6 RMSD) maps is shown in green/red meshes. PEG (magenta) was assigned to the electron density detected near the outer edge of the YcjN binding pocket near tyrosine 95 and tyrosine 72. No other unassigned electron densities were detected in the binding pocket. B. Ligand binding pockets of tmMBP3, mtUgpB, xaMalE, TTHA0356, and sgGacH proteins. All proteins are shown in gray ribbon representations and the negative volume imprints of their binding pockets are displayed in different colors, as determined by CASTp 3.0.

**Figure S4:**
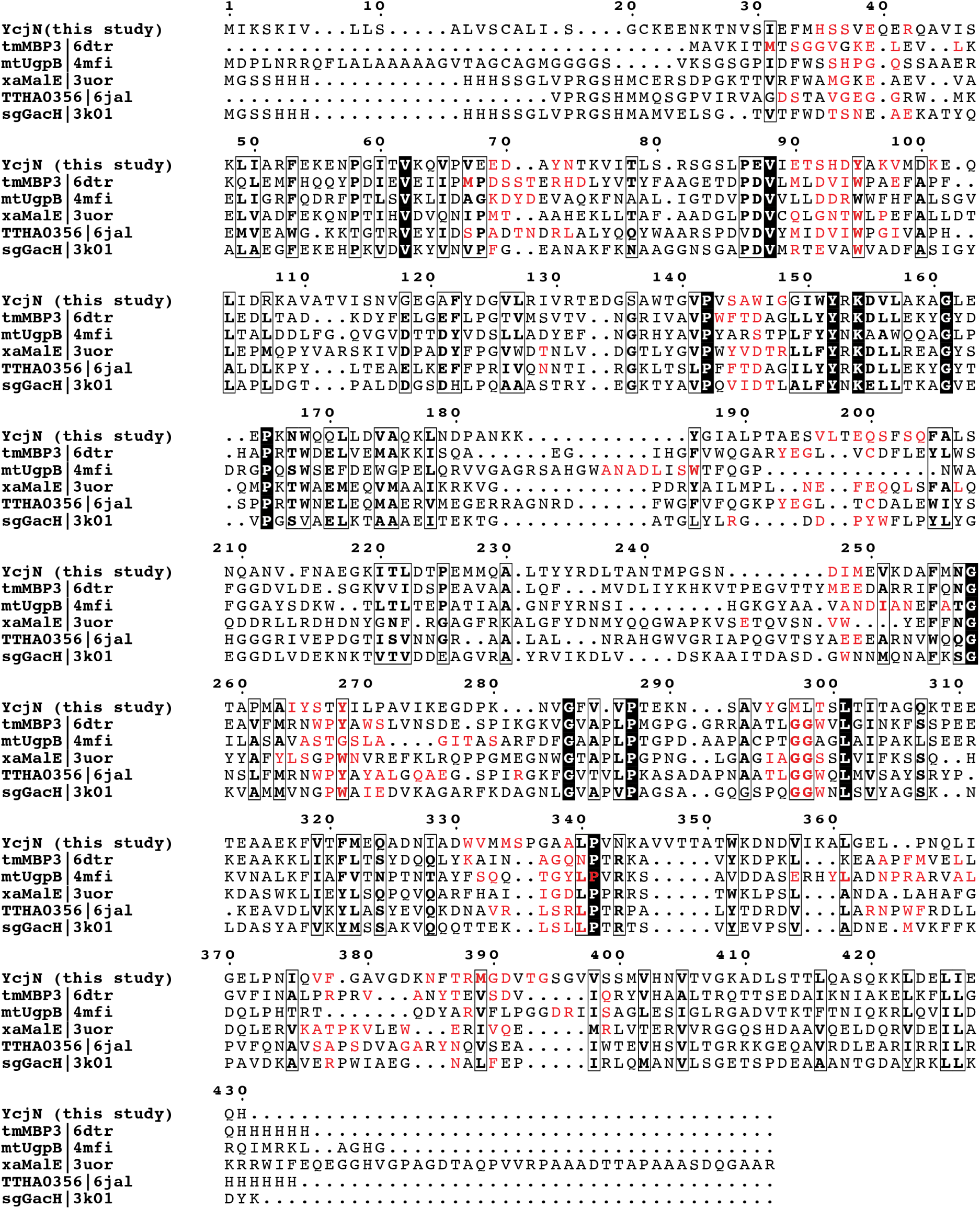
Sequence alignment of YcjN against top Dali hits. In the figure, white characters in a black box indicate strict identity, bolded characters indicate similarity within a group, a black frame indicates similarity across groups, and residues lining each protein’s binding pocket, as determined by CASTp 3.0, are colored in red.

